# High-throughput Tn-seq screens identify both known and novel *Pseudomonas putida* KT2440 genes involved in metal resistance

**DOI:** 10.1101/2024.04.12.589247

**Authors:** Kevin Royet, Laura Kergoat, Stefanie Lutz, Charlotte Oriol, Nicolas Parisot, Christian Schori, Christian H. Ahrens, Agnes Rodrigue, Erwan Gueguen

**Author notes:** These authors, in reason of their equal contribution to this work, have been placed in alphabetical order (Laura Kergoat, Stefanie Lutz, Charlotte Oriol, Nicolas Parisot, Christian Schori).

## Abstract

Chemical waste with toxic effects is released into the environment by industrial and urban activities. *Pseudomonas putida*, a rhizosphere bacterium, harbors a wide variety of genes capable of degrading hydrocarbons and xenobiotic compounds in its natural environment. This bacterium harbors also a large set of metal resistance genes. Most studies that identify genes involved in metal resistance in *P. putida* focus on over/underexpressed genes and may miss other genes important for metal resistance whose expression does not change. In this study, we used a Tn-seq approach to determine the essential genome of *P. putida* required for growth in the presence of an excess of metals in a culture medium. Tn-seq enables the detection of mutants with reduced or increased fitness in the presence of metal excess. We validated our screen by identifying known metal resistance gene such as *czcA-1* (*PP_0043*), *cadA-3* (*PP_5139*), *cadR* (*PP_5140*) and *pcoA2* (*PP_5380*). Their mutants were underrepresented in the presence of zinc, cadmium (for *cadA-3* and *cadR*) or copper respectively. In this study, we demonstrate by targeted mutagenesis and complementation assay that *PP_5337* and *PP_0887* are putative transcriptional regulators involved in copper and cadmium resistance, respectively, in *P. putida*. The study revealed the role of two genes, *PP_1663* and *PP_5002*, in cadmium and cobalt resistance respectively. This is the first evidence linking these genes to metal resistance and highlights the incomplete understanding of metal resistance mechanisms in *P. putida*.

## Introduction

*Pseudomonas putida* is an ubiquitous saprophytic bacterium that can utilize various sources of carbon and energy. This soil microorganism has been widely used as an experimental model to study the biodegradation of aromatic compounds or hydrocarbons (1, 2). It is able to colonize various habitats and has been isolated from water, soil, and the plant rhizosphere, sometimes polluted by various compounds (1, 3). The analysis of its complete genome revealed that *P. putida* possesses a wide range of genes that are involved in metal homeostasis or resistance. This suggests that the bacteria can survive in metal-polluted environments (4).

Metals play a crucial role in several cellular processes of microorganisms. Certain metals, such as nickel, cobalt, copper, and zinc, are essential nutrients. They function as stabilizers of protein structures or bacterial cell walls, as catalysts for biochemical reactions, and help maintain osmotic balance (5, 6). However, some metals can be highly toxic to cells. Metal toxicity can occur in various ways, such as oxidative damage caused by the production of reactive oxygen species, DNA damage, and protein damage due to the displacement of essential metals from their native binding sites or binding to respiratory enzymes (5). While some metals are essential for cellular function, they can also be toxic when present in excess. Therefore, it is crucial to tightly regulate the concentration of metal ions in cells to maintain optimal cellular activity. To maintain metal homeostasis, bacteria use various systems, such as metal uptake/efflux, chelation, or sequestration (5, 6). Several systems have been identified in *Pseudomonas putida* that confer resistance to heavy metals such as cadmium, zinc, and cobalt. These systems include P-type ATPases (e.g., CadA for cadmium resistance or CzcA for cadmium, zinc, and cobalt resistance), sequestration proteins (e.g., CopA for copper resistance), uptake systems (e.g., ZnuB/C or NiKABCDE), regulators, and proteins involved in redox mechanisms (4, 7, 8). Regarding metal stresses, a complete genome analysis has shown that the *P. putida* genome contains 61 open reading frames that are probably involved in metal tolerance or homeostasis, and seven more that are possibly involved in metal resistance (4). Proteomic or transcriptomic technologies have been used to investigate the response of *P. putida* to inhibitory concentrations of various metals (7–11). These studies indicated that a significant number of genes in *P. putida* are responsible for maintaining homeostasis, as well as tolerating and resisting various metals.

Although omics approaches (proteomics and transcriptomics) are considered powerful, they have limitations in terms of detectability and typically only detect genes whose expression levels significantly change between two conditions. These analyses may thus miss important factors not affected by gene/protein level changes. Therefore, more comprehensive screens are needed to identify new factors and ideally complete sets of genes involved in metal resistance in *P. putida*. The screening of *P. putida* CD2 mutants obtained by Tn5 insertions was also conducted in the presence of cadmium, which confirmed and completed the overview of the *P. putida* stress responses (12, 13). However, the low saturation levels of the Tn5 libraries suggests that some genes may have been missed during these analyses. Additionally, the number of tested mutants was limited by the need to test each individual mutant in every condition.

To gain a more comprehensive understanding of the genes necessary for metal resistance in *P. putida*, we utilized a high-throughput sequencing of a saturated transposon library (Tn-seq) (14, 15) in this study to screen tens of thousands of random insertion mutants of *P. putida* in the presence of excess amounts of metal ions. Tn-seq is a powerful method that has successfully been used to characterize essential genes in various conditions and many different species. For example, it has been used to identify essential genes for human gut colonization of B*acteroides thetaiotaomicron* (16), mouse colonization by human pathogens such as *Vibrio cholerae*, *Pseudomonas aeruginosa*, or *Streptococcus pneumoniae* (14, 17, 18), plant colonization by phytopathogens (19–21), tobramycin resistance genes of *P. aeruginosa* (22), toxic compound resistance genes in *P. putida* (23) and identification of genetic targets for improved tolerance of *P. putida* towards compounds relevant to lignin conversion (24). This technique has also been also employed to identify gold, silver and copper resistance genes in *Burkholderia cenocepacia* (25, 26). However, this technology has not yet been employed to discover genes involved in metal resistance in *P. putida* KT2440. By applying Tn-seq to screen a *P. putida* KT2440 mutant library in the presence of metals in the culture medium, we identified numerous genes required for growth in a culture medium rich in cobalt, copper, zinc (essential metals), or cadmium (a non-essential metal). Among them were *czcA-1* (*PP_0043*), *cadA-3* (*PP_5139*), *cadR* (*PP_5140*) and *pcoA2* (*PP_5380*), which are already known to be involved in zinc, cadmium, and copper resistance, respectively, and thereby validating the approach employed. In addition, we discovered several genes that were previously not associated with metal resistance and validated them through in-frame deletion and complementation assays. Our findings demonstrate that *PP_1663* (Cd), *roxSR* (Cd), *PP_5337* (Cu), and PP_5002 (Co) all are involved in metal resistance in *P. putida*.

## Results and discussion

### Characterization of *P. putida* KT2440 Himar1 transposon library

Tn-seq screening has been performed numerous times with the opportunistic pathogen *Pseudomonas aeruginosa*. An elegant strategy that has been developed involves the use of a modified *Himar9* mariner transposon derivative carrying Mme1 restriction sites in the inverted repeats (IR) and a gentamicin resistance cassette between the IRs (18). The Mariner transposon can specifically insert itself into the genome at TA sites. In *P. putida* KT2440, 129,002 TA sites can be targeted by this transposon. To generate a pool of approximately 1,000,000 colonies, we introduced by conjugation from *E. coli* the plasposon pSam_D-Gm into *P. putida* KT2440 (18). Two technical replicates of the DNA libraries were created from this pool and were subjected to high-throughput sequencing. The TPP software (27) was used to determine the number of reads at each TA site. The sequencing of the two samples detected 91,882 and 93,147 unique insertions into TA sites, with an average of 91 and 96 reads per TA, respectively (Table S3). The preparation of the Tn-seq library was highly reproducible, with a Pearson correlation coefficient of 98%. The density of Tn insertions was approximately 70% from our initial pool of mutants (Table S3), and the unique insertions were distributed all around the chromosome (Fig. S1). These results indicate high quality and coverage of our *P. putida* Tn-seq libraries.

The gene essentiality of the Tn-seq input libraries was next determined using the TRANSIT software (27), which employs a Hidden Markov Model (HMM) method to predict essentiality and non-essentiality for individual insertion sites (DeJesus & Ioerger 2013). The HMM analysis identified 600 genes essential for growth on LB agar, representing 10.8% of the genes of *P. putida* KT2440. 4458 genes were identified as non-essential genes (NE) (Table S4).

### Screening of genes important for metal resistance

To identify new genes responsible for metal resistance in *P. putida* KT2440, we tested the effects of copper chloride, zinc, cobalt, and cadmium on the growth of *P. putida*. We used LB rich medium in our screens instead of minimal medium to prevent the loss of auxotrophic mutants or biosynthesis pathways that could be important for metal resistance during growth. To determine the optimal metal concentration, we grew *P. putida* KT2440 in LB rich medium supplemented with varying concentrations of a metal ion solution. We compared the growth of the WT strain with and without excess Cu, Zn, Cd, or Co under the same conditions used for screening, i.e., in an Erlenmeyer flask with shaking at 30°C (Fig. S2). Under our laboratory conditions, we found that a concentration of 10 µM cobalt chloride, 2.5 mM copper chloride, 125 µM zinc chloride, and 12.5 µM cadmium chloride did not affect the growth of the cultures during the exponential phase, compared to the growth of the WT strain grown without an excess of these metals (Fig. S2). We hypothesized that under these conditions, only homeostatic mechanisms will be selectively activated, rather than pleiotropic responses to metal toxicity.

For the Tn-seq screening, biological replicates were performed to ensure the reproducibility of the method. The cultures were inoculated with 10^7^ bacteria from the mutant pool. After twelve divisions in the presence of metals in the culture medium, at 28°C, the final pools of mutants were collected. Sequencing of transposon insertion sites of the final pools, followed by the TPP analysis, indicated highly reproducible results with a Pearson correlation coefficient > 90% for each dataset (Table S3).

To test the statistical significance of the genes that contribute to *P. putida*’s loss or gain of fitness in a metal-rich medium, we conducted a RESAMPLING (permutation test) analysis using the TRANSIT software. We compared the results obtained from culture in LB to those obtained from culture in LB with an excess of metal (Table S5). After applying Tn-seq to our datasets and selecting only genes with an FDR adjusted p-value (q-value) ≤ 0.05, we identified 9 genes involved in cobalt resistance, 14 in copper resistance, 3 in zinc resistance, and 8 in cadmium resistance. From these 28 genes, we applied an additional cutoff by removing 3 genes with a mean read count in LB below 2 (less than 2 reads on average per TA) and that are classified as essential or causing growth defects in LB. Finally, we retained 25 genes (Table 1). 23 genes were classified as non-essential in LB, while the remaining 2 were identified as causing growth defects and growth advantages. Some of these genes, highlighted in bold, were previously known to be involved in metal resistance in *P. putida*, thus confirming the validity of the Tn-seq approach. In the following sections, we discuss the function of some of the genes we consider most important in relation to metal resistance.

**Table 1.** Metal resistance genes of *P. putida* KT2440 discovered by Tn-seq.

| Metal<sup>a</sup> | Locus | Gene<sup>b</sup> | Function | State in LB<sup>c</sup> | No. of TAs | Mean read<sup>d</sup> |  | Log<sub>2</sub>FC<sup>e</sup> | q-value<sup>f</sup> | orthologs in PAO1<sup>g</sup> |
| --- | --- | --- | --- | --- | --- | --- | --- | --- | --- | --- |
|  |  |  |  |  |  | LB | LB + metal |  |  |  |
| Co | PP_2645 | mgtA | Magnesium transporter ATP-dependent | GD | 24 | 6.1 | 78.9 | 3.70 | 0.00000 |  |
|  | PP_5002 | PP_5002 | Hypothetical protein | NE | 5 | 38.1 | 0.2 | -7.31 | 0.00000 | PA5055 |
|  | PP_0096 | prlC | Oligopeptidase A | NE | 30 | 126.0 | 23.4 | -2.43 | 0.00000 |  |
|  | PP_0340 | glnE | Glutamate-ammonia-ligase adenylyltransferase | NE | 37 | 28.1 | 6.5 | -2.12 | 0.00000 |  |
|  | PP_5328 | pstC | Phosphate ABC transporter permease | NE | 32 | 33.6 | 8.7 | -1.95 | 0.00000 |  |
|  | PP_0243 | gshA | Glutamate-cysteine ligase | NE | 28 | 16.5 | 5.6 | -1.56 | 0.00000 |  |
| Cu | PP_0691 | proB | Glutamate 5-kinase | NE | 7 | 3.8 | 55.8 | 3.89 | 0.00000 |  |
|  | **PP_0586** | **cadA2 → copA1** | **Cadmium translocating P-type ATPase** | NE | 22 | 197.3 | 0.7 | -8.19 | 0.00000 | PA3920 = copA1 |
|  | PP_1735 | htrB | Lipid A biosynthesis lauroyl acyltransferase | NE | 10 | 23.6 | 0.1 | -7.39 | 0.00000 |  |
|  | PP_4216 | pvdE | Pyoverdine ABC transporter ATP-binding protein/permease | NE | 18 | 49.0 | 1.1 | -5.45 | 0.00000 |  |
|  | PP_4213 | pvdM | Dipeptidase | NE | 15 | 58.7 | 2.9 | -4.35 | 0.00000 |  |
|  | PP_4214 | pvdN | Pyoverdine biosynthesis-like protein | NE | 15 | 62.4 | 2.8 | -4.49 | 0.00000 |  |
|  | PP_4215 | pvdO | Pyoverdine biosynthesis-like protein | NE | 18 | 72.6 | 4.0 | -4.18 | 0.00000 |  |
|  | PP_2767 | PP_2767 | ABC transporter ATP-binding protein | NE | 5 | 22.1 | 1.6 | -3.81 | 0.00000 |  |
|  | PP_5337 | PP_5337 | LysR family transcriptional regulator | NE | 11 | 19.5 | 2.4 | -3.04 | 0.00000 | PA5428 |
|  | **PP_5380** | **copA-2 → pcoA2** | **copper resistance protein A** | NE | 36 | 259.6 | 32.3 | -3.01 | 0.00000 | PA2065 = pcoA |
|  | **PP_5379** | **copB-2 → pcoB2** | **copper resistance protein B** | NE | 17 | 113.1 | 33.5 | -1.75 | 0.00000 | PA2064 = pcoB |
|  | PP_4194 | gltA | Citrate synthase | NE | 21 | 30.8 | 10.0 | -1.62 | 0.00000 |  |
|  | PP_0243 | gshA | Glutamate-cysteine ligase | NE | 28 | 16.5 | 5.9 | -1.50 | 0.00000 |  |
|  | PP_2328 | cysH | Phosphoadenosine phosphosulfate reductase | GA | 11 | 390.5 | 158.9 | -1.30 | 0.00000 |  |
| Zn | **PP_5140** | **PP_5140 → cadR** | **MerR family transcriptional regulator** | NE | 9 | 181.6 | 9.1 | -4.31 | 0.00000 | PA3689 |
|  | PP_4213 | pvdM | Dipeptidase | NE | 15 | 58.7 | 18.4 | -1.67 | 0.00000 |  |
|  | **PP_0043** | **czcA-1** | **Cation efflux system protein** | NE | 51 | 211.3 | 90.6 | -1.22 | 0.00000 | PA2520 = czcA |
| Cd | PP_1663 | PP_1663 | Hypothetical protein | NE | 10 | 364.6 | 1.2 | -8.22 | 0.00000 | PA0943 |
|  | PP_0127 | dsbA | Thiol:disulfide interchange protein | NE | 8 | 32.2 | 0.2 | -7.59 | 0.00000 |  |
|  | PP_5140 | PP_5140 → cadR | MerR family transcriptional regulator | NE | 9 | 181.6 | 1.7 | -6.74 | 0.00000 |  |
|  | **PP_5139** | **cadA-3** | **cadmium translocating P-type ATPase** | NE | 18 | 110.4 | 3.5 | -4.99 | 0.00000 |  |
|  | PP_4216 | pvdE | Pyoverdine ABC transporter ATP-binding protein/permease | NE | 18 | 49.0 | 4.1 | -3.59 | 0.00000 |  |
|  | PP_0887 | PP_0887 → roxS | Sensor histidine kinase | NE | 19 | 193.2 | 19.5 | -3.31 | 0.00000 | PA4494 = RoxS |
|  | PP_4213 | pvdM | Dipeptidase | NE | 15 | 58.7 | 7.4 | -2.98 | 0.00000 |  |
|  | PP_4214 | pvdN | Pyoverdine biosynthesis-like protein | NE | 15 | 62.4 | 8.6 | -2.86 | 0.00000 |  |

<sup>a</sup> Metal tested for which a significant log<sub>2</sub>FC has been calculated.
<sup>b</sup> Name of the gene in the TIGR KT2440 genome of the *Pseudomonas* database (version 22.1, date: 2023-10-06). When named has evolved in other *Pseudomonas* species, new names of the genes have been indicated in order to align with homologies.
<sup>c</sup> State of each gene in LB defined by the TRANSIT software using an Hidden Markov Model: NE, Non-Essential ; GD, Growth-Defect ; E, Essential ; GA, Growth-Advantage.
<sup>d</sup> Mean reads per TA site for a gene in each growth condition.
<sup>e</sup> Ratio of reads between the two conditions expressed in log<sub>2</sub>.
<sup>f</sup> P-values adjusted for multiple comparisons using the Benjamini-Hochberg procedure (See Transit manual).
<sup>g</sup> Orthologous gene in *Pseudomonas aeruginosa* PAO1.

### Analysis of the genes of *P. putida* required for metal resistance

#### Copper resistance

Genes required for copper resistance were identified using a subinhibitory concentration of copper (see Figure S2). Copper is an essential metal required as a cofactor for electron transport and redox enzyme systems in aerobic bacteria. In bacteria, copper exists in two different states: the less toxic oxidized Cu(II) state, which can be transformed into the more toxic reduced Cu(I) state under redox systems. Copper is toxic to cells because it can displace other metals from essential complexes and bind to various biomolecules in an unspecific manner. Fourteen candidate genes, listed in Table 1, were found to be possibly involved in copper resistance. The function of these genes is discussed below.

##### Inner membrane protein Cu^+^-ATPase

The role of *P_1B_-type ATPases* in copper resistance has been extensively studied in *P. aeruginosa* PAO1. The transmembrane inner membrane protein P_1B_-type ATPase, CopA, is responsible for cytoplasmic Cu^+^ efflux. *P. aeruginosa* PAO1 has two homologous Cu^+^-ATPases, CopA1_PAO1_ (PA3920) and CopA2_PAO1_ (PA1549). CopA1_PAO1_ was expressed in response to high Cu^+^ (28–30), and its deletion induced copper sensitivity (31). However, while CopA2_PAO1_ does not directly contribute to copper resistance, it does play a crucial role in loading copper into cytochrome c oxidase subunits. Both enzymes export cytoplasmic Cu^+^ into the periplasm (31). In *P. putida* KT2440, only one copper Cu^+^-ATPase is present, which is encoded by *PP_0586*. The protein is commonly referred to as CadA2 in *P. putida* KT2440. However, we will use the name CopA1_KT2440_ in the further section in order to align with naming and homologies, since *copA1*_KT2440_ is the orthologous gene of CopA1_PAO1_.To ensure that the two proteins were the same, we compared their 3D structures predicted by AlphaFold (32) using the TM-Align algorithm (33). The structures were found to be superimposed (Fig. S3). Previous research has demonstrated that CopA1_KT2440_ is highly produced in the presence of copper in minimal salt media (7). Our Tn-seq screen in the presence of copper revealed that *copA1*_KT2440_ mutants have a strong growth disadvantage (Table 1). In *P. aeruginosa*, *copA1_PAO1_* is positively regulated by CueR (PA4778) (28–30). The transcriptional regulation of *copA1*_KT2440_ by CueR_KT2440_ has not been verified in *P. putida.* however, a putative cueR binding site (ACCTTGCCTGCGTGGCAAGGT) is located in the promoter region of *copA1*_KT2440_ as indicated in the RegPrecise database (https://regprecise.lbl.gov/sites.jsp?regulog_id=5159 ; (34)), suggesting a direct regulation like in P. aeruginosa.

##### Outer membrane protein PcoB and putative periplasmic multi-copper oxidase PcoA

In certain bacterial species, an outer membrane porin called PcoB appears to contribute to periplasmic Cu^+^ efflux. *pcoB* is often co-localized with *pcoA* encoding a putative periplasmic multi-copper oxidase (35). *pcoAB* were first identified as part of a copper resistance Cop operon in the pPT23D plasmid of *Pseudomonas syringae* (36). For this reason, PcoA proteins were sometimes mistakenly named CopA, despite being functionally distinct from the previously described CopA proteins which are ATPases of the inner membrane (7, 36). In *P. syringae*, PcoA and PcoB bind copper (36). In *P. aeruginosa*, the orthologous system named *pcoA/B* (*PA2065* and *PA2064*) is induced up to 70-fold in the presence of a high copper sulfate concentration (28, 30). In *P. putida* KT2440, the orthologous genes of *PA2065* and *PA2064* are *PP_5379* (*copB-2*) and *PP_5380* (*copA2*) respectively. The AlphaFold predicted 3D structure alignment of the orthologous proteins using the TM-Align algorithm confirms that these proteins adopt the same conformation (Fig. S3). For clarity, we thus decided to rename these P. putida KT2440 genes *pcoB-2* and *pcoA-2* respectively. Mutants of these genes exhibit growth-defect phenotypes in the presence of copper (Table 1 and Fig 1). *P. putida* KT2440 also has a second PcoAB system (pcoA-1/pcoB-1) encoded by *PP_2204*/*PP_2205*, but mutants of these genes do not exhibit any growth-defects in the presence of copper. As only the PcoA-2/B-2 system appears to be essential for copper resistance, it is possible that the PcoA-1/B-1 system is not expressed under our laboratory conditions or is less efficient than the PcoA-2/B-2 system.

**Figure 1.**
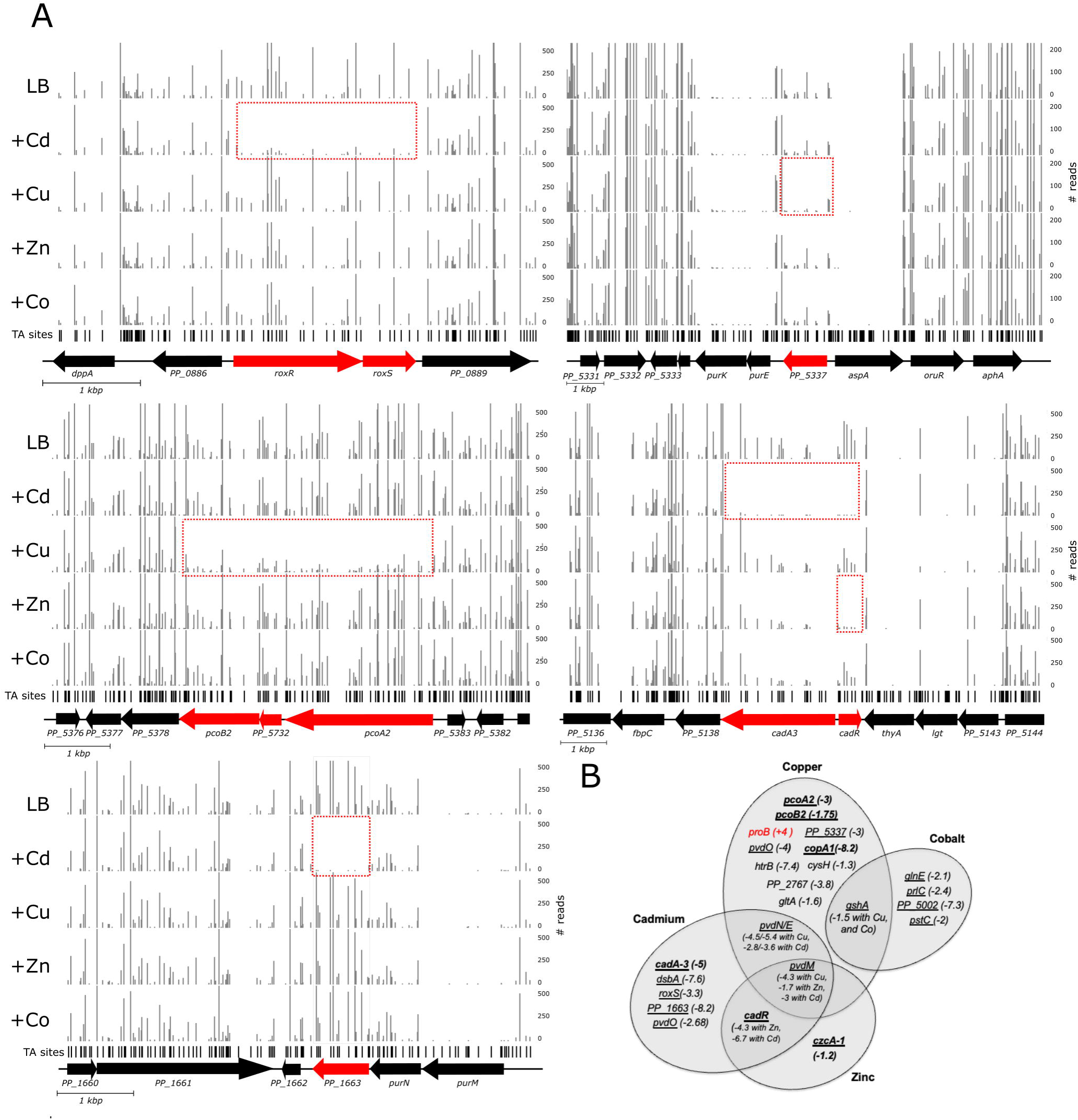
Genes involved in metal resistance according to the Tn-seq experiment. (A) Examples of negative selection revealed by Tn-seq. The graphs show the number of Tn-seq reads at each location aligned to TA sites on the *P. putida* KT2440 genome. Results are shown in LB only or LB with either Cu, Cd, Zn or Co. The regions with significantly fewer reads are framed in red and the genes corresponding to these regions are indicated in red. Data are averaged from biological replicates and normalized as described in Materials and Methods. (B) Venn diagram of the genes with a positive or negative log_2_FC, indicating the fitness difference between the test condition (LB with a excess of Cu, Cd, Co or Zn) and the LB condition. Genes already known to be involved in metal resistance in *P. putida* are in bold. The underlined genes were selected for in-frame deletion and further analysis.

*A*fter copper treatment, *pcoA-2* and *pcoB-2* are highly transcribed in *P. putida* KT2440, and PcoB-2 accumulates in cells (7). In some strains of *E. coli* that harbor an episomal gene cluster *pcoABCDRSE*, the Pco operon is mediated by the PcoRS two-component system. In *P. aeruginosa* PAO1, PcoA/B expression is suppressed in the ΔcopR strain (Teitzel et al. 2006; Miller et al. 2009; Quintana et al. 2017). In *P. putida* KT2440, we searched for the consensus CopR binding site TGACANNNNTGTNAT (30) and found it in the intergenic region upstream of *copRI/CopS1* (*PP_2158/PP_2157*) and *pcoA-1/pcoB-1* (*PP_2205/PP_2204*). We did not detect any putative CopR binding site within the promoter region of *pcoA2/B2,* suggesting that these genes may be activated differently under copper stress. *P. putida* has two CopR regulators, CopR1 (PP_2158) and CopR2 (PP_5383). The proximity of *copR2* to *pcoA-2/B-2* suggest that CopR2 might regulate *pcoA2/B2*. However, this hypothesis has not yet been investigated.

##### The Cus system

In *P. putida* KT2440, the genes *PP_5379* (*pcoB-2*) and *PP_5380* (*pcoA-2*) are located adjacent to a cluster of five genes, *copR1/S1* and *cusCBAF*. *cusCBA* genes encode a putative cation/proton antiporter that spans the outer and inner membranes and has been proposed to be involved in copper and silver efflux (4). The two-component system CusR/S senses periplasmic copper and regulates the Cus RND-type transport system in *E. coli* (37, 38). However, our Tn-seq datasets did not reveal any potential role of the CusCBAF system and CopRS two-component regulatory systems in copper resistance at the copper concentration we used. This confirms an earlier observation showing that *cusC* expression is not activated in the presence of copper (7). The *cusCBAF* operon, which was suggested to be involved in copper resistance, may respond to a different concentration of copper.

#### Cadmium resistance

Cadmium commonly forms cations with an oxidation state of II. Unlike zinc, cadmium has a preference for binding to sulfur ligands, which can be problematic for proteins with disulfide bonds. Zinc homeostasis and cadmium resistance mechanisms often overlap due to their similarities. They share uptake and efflux transporters, as well as metal-responsive regulatory proteins (4, 7, 39). Cadmium can be removed from the cytoplasm of bacterial cells through various systems, including the P-type ATPase CadA-3 (PP_5139). CadA-3 is a homolog of ZntA in *E. coli* (4, 40, 41). Our Tn-seq screening indicates that CadA-3 is involved in cadmium resistance, with a strong negative log_2_FC of −4.99 (table 1 and fig. 1). It confirms previous observations that CadA-3 confers resistance to Cd^2+^, while CadA1 plays no role in resistance to Cd, Zn, Cu, or Co in *P. putida* (39). The orthologous gene of *cadA-3* in *P. putida* CD2 was previously shown to be a major determinant of cadmium resistance (12).

The *dsbA* gene, which is essential for cadmium resistance, was discovered with a negative log_2_FC of −7.59 (Table 1 and Fig. 1). DsbA catalyzes the oxidation of disulfide bonds of periplasmic proteins. As a result, the cysteine residues of DsbA become reduced, and the protein must be oxidized by DsbB to be regenerated (42). Notably, a *dsbA* mutant was found to be sensitive to cadmium and even zinc in both *E. coli* (43) and *Burkholderia cepacia* (44). In contrast, a *dsbB* mutant did not appear to be essential for cadmium resistance under the test conditions.

Our Tn-seq analysis showed that the gene encoding the RoxS sensor (PP_0887), which is part of the RoxS-RoxR two-component system (PP_0887-PP_0888), is essential for cadmium resistance. The two genes coding this two-component system are transcribed in a single unit (45). The system belongs to the RegA/RegB family, where RegA functions as an integral membrane sensor histidine kinase, and RegB is a sigma 54-dependent regulator. A whole-genome transcriptional analysis was conducted to define the *P. putida* RoxS/RoxR regulon in LB. The regulon includes genes involved in amino acid and sugar metabolism, the sulfur starvation response, elements of the respiratory chain, and genes that participate in maintaining the redox balance (45). Although a putative RoxR recognition element has been identified in the promoters of genes regulated by this system (45), the specific genes that are up or down regulated by RoxS/RoxR in response to cadmium are still unknown.

#### Cobalt resistance

Cobalt is a transition metal with an oxidation state of II. It plays an essential role for microorganisms as cofactors for diverse metalloenzymes. Cobalt toxicity is related to its potential interference with iron and possibly manganese homeostasis. Bacteria typically use efflux systems to survive in an environment with an excess of Co^2+^. The cobalt resistance system was poorly described in *Pseudomonas*, but it was studied in greater detail in other organisms (46).

The genes of *P. putida* KT2440 involved in cobalt resistance were determined using a sub-inhibitory concentration of cobalt. Our screen did not reveal any systems that cause cobalt resistance in other bacteria and that exist in *P. putida*. The *czcCBA* RND system in *Cupriavidus metallidurans* confers resistance to Cd^2+^, Zn^2+^, and Co^2+^ (47). Although the CzcCBA system exists in KT2440, it was only reported to confer Zn^2+^ and Cd^2+^ resistance (39). Additionally, it was discovered that CzcD, a member of the CDF family, confers cobalt resistance in *Ralstonia sp*. Strain CH34, although to a lesser extent than the CzcCBA system (48). A homolog of *czcD*, *PP_0026*, exist in *P. putida* KT2440, but it does not confer cobalt resistance in our Tn-seq screening. Interestingly, our screen did not reveal a role of the MrdH efflux pump (PP_2968), which is homologous to the RcnA efflux pump from *E. coli*. Although cobalt induces *mrdH* activity, the efflux pump does not confer resistance to cobalt (49).

Although our screening did not identify any genes encoding efflux pumps, we did identify some genes that were not previously known to be involved in cobalt homeostasis. One of these genes is mgtA, which encodes an ATP-dependent magnesium transporter that is involved in the active transport of magnesium in cells. This gene has a positive log_2_FC value, indicating that the mutant confers a growth advantage in the presence of cobalt in the culture medium. Although MgtA has not been experimentally characterized in *P. putida* or *E. coli*, its ortholog in *Salmonella typhimurium* has been studied, where MgtA mediates magnesium uptake (50, 51).

#### Zinc resistance

Zinc has an affinity for ligands containing oxygen, nitrogen, or sulfur and is often used as an enzyme cofactor in the cell. As mentioned for copper, zinc toxicity occurs with its ability to replace another metal from enzymes or by forming complexes with other biomolecules. It exists in cells mainly in the oxidized state Zn^2+^. Zinc homeostasis is well documented and is regulated by several processes: Zn^2+^ uptake regulation, sequestration by metallothioneins (MT) and efflux system (52, 53). Our screen revealed three genes involved in zinc resistance: *czcA-1* (*PP_0043*), *cadR* (*PP_5140*) and *pvdM* (*PP_4213*). The CzcCBA system has been fully described in bacteria. It has been reported to confer Cd^2+^, Zn^2+^ and Co^2+^ resistance in *C. metallidurans* and Cd^2+^ and Zn^2+^ resistance in *P. putida* KT2440 (39, 46). The identification of the main component of the CzcCBA system, CzcA-1, confirms the validity of our screening in the presence of zinc. The CzcCBA system may be predominant at the zinc concentration used. At least five *czcA* genes have been described in *P. putida* KT2440 (4). Our screen confirms a previous result showing that CzcCBA1 is the predominant CzcCBA system in *P. putida* under laboratory conditions (39).

#### Cross metal resistance

##### Pyoverdine

Pyoverdine is the major siderophore in fluorescent *Pseudomonads*. The pyoverdine pathway is complex, with 20 different proteins documented to be involved in its regulation, synthesis, maturation, transport and uptake (54). Pyoverdine maturation starts with the transport of a precursor (PVDIq) from the cytoplasm to the periplasm by the ABC transporter PvdE. PvdN and PvdO are involved in the maturation of the pyoverdine precursor (55, 56). PvdM is required for the oxidation of ferribactin by PvdP during periplasmic pyoverdine maturation (57). The p*vdM,N,O,E* genes belong to an operon in *P. putida* KT2440 but not in *P. aeruginosa* PAO1. The mature pyoverdine is able to chelate many metals, but with a lower affinity than iron (Schalk & Guillon 2013), and could thus protect the cell from metal toxicity (54, 58). In our screen, since all mutants grow in the same culture medium, a mutant defective in the production of pyoverdine can be protected by the pyoverdine produced by the other mutants. However, at least one gene of the *pvdMNOE* operon was found to be involved in copper, cadmium or zinc resistance (Fig. 1 and Table 1). Although not statistically significant for cadmium resistance, the *pvdO* gene has a log_2_FC of −2.68, similar to the other genes in the operon. Similarly, the *pvdN,O,E* genes have a log_2_FC of −1.56, −1.26 and −2.01, respectively, in the presence of zinc. The whole *pvdMNOE* operon seems to be important for copper, zinc and cadmium resistance (Fig. 1, Table 1). This is consistent with the proposed hypothesis that mature periplasmic storage of pyoverdine protects the bacterium from excess metals other than iron by chelating these metals in the periplasm (54).

##### gshA

Among the genes involved in copper and zinc resistance, *gshA* (PP_0243) was found in our Tn-seq screen. *gshA* encodes the glutamate cysteine ligase GshA, which forms the glutamyl-cysteine from L-glutamate and is essential for copper resistance with a log_2_FC of −1.5 (Fig. 1 and Table 1). Glutamyl-cysteine is itself used by GshB to produce glutathione. Glutathione is a key player in metal homeostasis in *E. coli* (59) and glutathione can buffer an excess of intracellular copper in *Streptococcus pyogenes* (60). The thiol group and cysteine residues of glutathione can directly bind to metal ions, protecting the cells from their deleterious properties. *gshB* (PP_4993) had a negative log_2_FC but did not pass the statistical threshold for copper resistance (log_2_FC of −1.04) (table S5). It is noteworthy that a mutant of the *proB* gene (PP_0691), involved in proline biosynthesis, provides a growth advantage in the presence of copper. *proB* encodes glutamate 5 kinase, which transforms L-glutamate into L-proline. However, L-glutamate is also the substrate of the glutamate cysteine ligase, which is produced by *gshA*. As the *gshA* gene appears to be essential for copper resistance, it is not surprising that a *proB* mutant confers a growth advantage in presence of copper. Likewise, it is not unexpected to detect *gshA* involved in a cross resistance since it has already been described for copper (II), zinc (II) and cadmium (II) resistance in *E. coli* (Helbig et al. 2008). However, it is the first time that *gshA* is described as being involved in cobalt resistance (59).

##### CadR

We also highlight the importance in metal resistance of the merR regulator CadR in metal (*PP_5140*), which has a strong negative log_2_FC of −6.74 for cadmium resistance and of −4.31 for zinc resistance (table 1 and fig. 1). *cadR* is the neighbor gene of cadA-3 (*PP_5139*), which was also implicated in cobalt resistance but not zinc resistance in our screen (see upper section). CadR regulates its own transcription and is known to respond to cadmium (61). According to Canovas and colleagues, CadR was described as the putative regulator of *cadA-3* (*PP_5139*), but this regulator does not regulate *cadA* in *P. putida* 06909 (61). Previous mutational analysis indicated that *cadA-3* and *cadR* are partially responsible for zinc resistance in *P. putida* 06909 (61). Although CadR preferentially binds to cadmium, it can also weakly bind to zinc, resulting in less transcription activation compared to when it is complexed with cadmium (62). In *P. aeruginosa*, CadR is constitutively bound to its promoter and promptly activating *cadA* gene expression upon Zn binding. CadA is essential for a timely induction of the CzcCBA efflux system (63). In our condition of an excess of zinc ions, cadA-3 was not required for zinc resistance.

#### In-frame deletion mutants and complementation assays confirmed that *PP_1663* and *roxSR* are required for Cd^2+^, *PP_5337* for Cu^2+^ and *PP_5002* for Co^2+^ are required for metal resistance

Our Tn-seq screen also identified several genes that were not previously known to be associated with metal resistance. Among them, PP_1663, a gene encoding a putative periplasmic protein of unknown function, was found to be involved in cadmium resistance with the strongest log_2_FC of −8.22. This gene is the ortholog of *PA0943* in *P. aeruginosa* PAO1. A *PA0943* mutation rendered *P. aeruginosa* hypersensitive to the production of the secretin XcpQ and altered the normal functioning of the Xcp protein export system (64). In *P. putida* KT2440, The type II secretion system (Xcp) of *P .putida* is involved in the secretion of phosphatase (65). We could also identify a new transcriptional regulator of the LysR family, *PP_5337*, probably involved in copper resistance (log_2_FC of −3.04). Analysis of its regulon has not been performed yet. The Tn-seq screen in presence of cobalt also identified the *PP_5002* gene with unknown function. The putative protein produced by *PP_5002* contains a DUF971 domain which could be involved in Fe-S cluster assembly. The *PP_5002* product could play a major role in iron homeostasis to counteract the deleterious effect of cobalt on this equilibrium. Finally, we identified the RoxS sensor (PP_0887), which is part of the RoxS-RoxR two-component system (PP_0887-PP_0888), as being essential for cadmium resistance.

To confirm the role of these genes in metal resistance, we decided to go a step further by performing in-frame deletions of these genes and selected other genes identified in our Tn-seq screening. We also made mutants of *pcoA-2/B-2* for copper resistance, *czcA-1* for zinc resistance, and *cadA-3* for cadmium resistance because they can be used as positive controls. Since *roxS* (*PP_0887*) is in operon with *roxR* (*PP_0888*), a double mutant was constructed. As several genes of the *pvdMNOE* operon were identified in our screening, we decided to make the Δ*pvdMNOE* mutant. The genes that were deleted are underlined in Figure 1B.

First, to validate the Tn-seq results, we performed a competition experiment with the WT strain and the mutants in a 1:1 ratio to calculate the fitness of the respective mutants compared to the WT strain. We calculated a ratio in log_10_ by dividing the number of colony-forming units (cfu) of the mutant by the number of cfu of the WT strain after a co-culture of the two strains. The experiment were performed in LB only or LB with a respective metal at the identical concentration that was used in the Tn-seq screen. (Fig. 2). First, In LB only, all tested mutants grew as well as the WT strain, except for the Δ*dsbA* and Δ*gshA* mutants, which showed reduced growth fitness. In the presence of cadmium, we confirmed that the Δ*roxSR,* Δ*pvdMNOE,* Δ*cadR*, Δ*dsbA,* Δ*PP_1663,* and Δ*cadA-3* mutants had a lower fitness than the WT strain (Fig. 2A). The growth of the Δ*dsbA* and Δ*cadA-3* mutants in the presence of cadmium was so low that the fitness could not be calculated. In the presence of copper, the Δ*pvdMNOE,* Δ*copA1,* Δ*gshA,* Δ*pcoA2,* Δ*pcoB2*, and Δ*PP_5337* mutants had a lower fitness (Fig. 2B). When exposed to cobalt, only the Δ*PP_5002* mutant showed a significant fitness defect. We were unable to confirm the sensitivity towards cobalt for the Δ*pstC*, Δ*gshA*, Δ*glnE*, and Δ*prlC* mutants (Fig. 2C). Finally, the mutants Δ*cadR*, Δ*pvdMNOE*, and Δ*czcA-1* exhibited lower fitness levels in the presence of zinc. However, the cadA-3 mutant showed no sensitivity to zinc (Fig. 2D). This confirms our Tn-seq results. In conclusion, we also showed that the Δ*cadR* mutant is sensitive to both cadmium and zinc and that the Δ*pvdMNOE* strain is sensitive to both cadmium, zinc and copper. In general, all results confirm our Tn-seq results, except for cobalt where only one gene (Δ*PP_5002*) could be validated.

**Figure 2.**
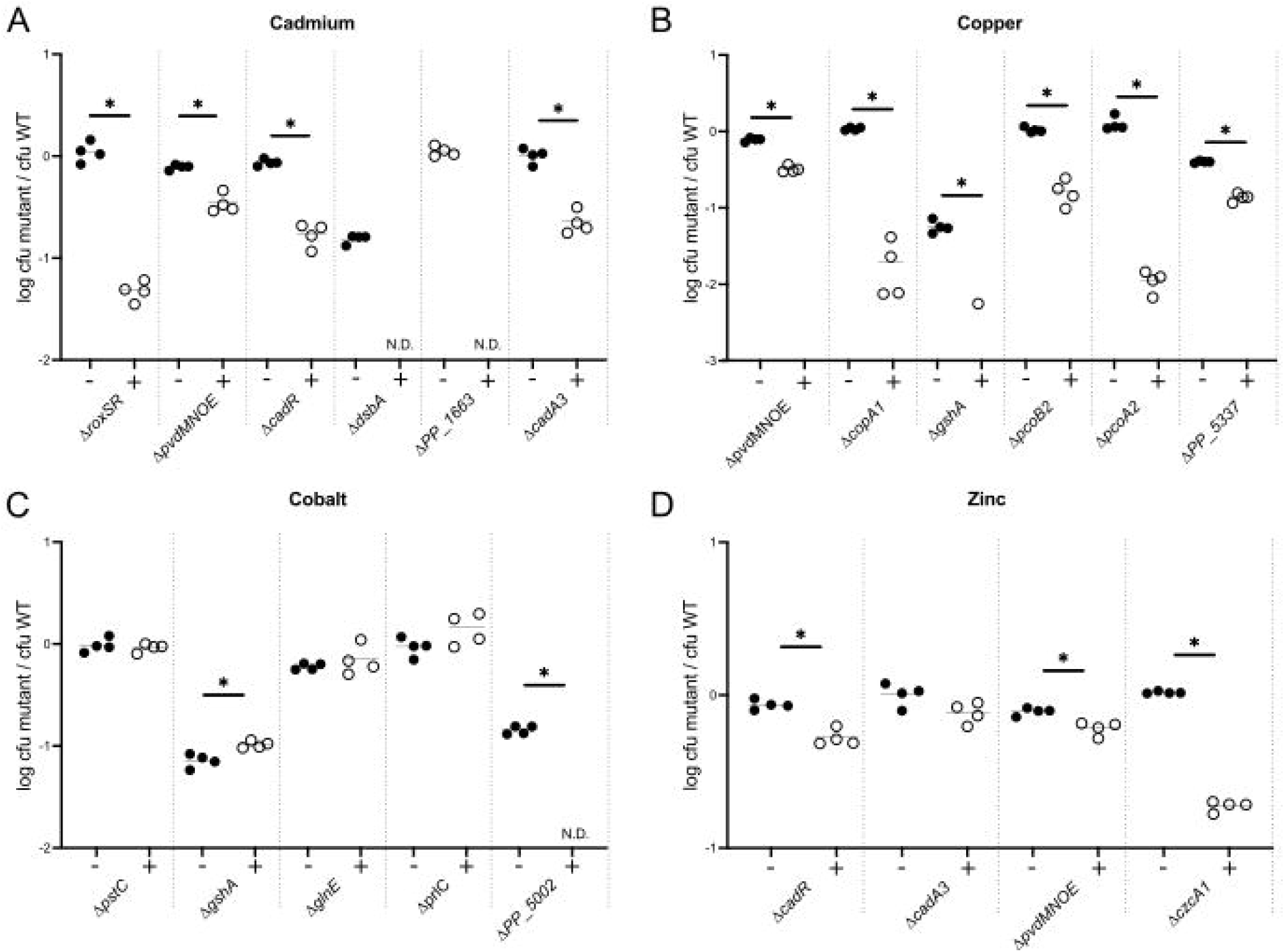
Competition between the wild type and the mutant strains of *P. putida* KT2440 in presence or not of a metal in excess. Competitions were realized with an initial ratio of 1:1 in LB supplemented or not with a sub-inhibitory concentration of metals (cobalt 10 µM, zinc 125 µM, copper 2.5 mM, cadmium 12.5 µM). The respective final ratio was determined as described in Methods and presented in Log_10_. The experiment was realized four times. * indicates a statistically significant difference relative to the absence of metal condition (p<0.05, Mann-Whitney U test).

Next, we focused our work on the four genes *PP_5337, roxSR, PP_1663* and *PP_5002* because they had not been shown to be involved in metal resistance prior to our work. The growth of these mutants was measured individually in LB liquid culture over time and compared to the growth of the WT strain (Fig. 3A-G). No statistical difference in growth was observed between the mutants and the WT strains. In contrast, in the presence of metal ions, the mutants Δ*PP_5337,* Δ*roxSR,* Δ*PP_1663* and Δ*PP_5002* showed a growth defect in LB supplemented with Cu, Cd, Cd and Co, respectively (Fig. 3A, C, E, G).

**Figure 3.**
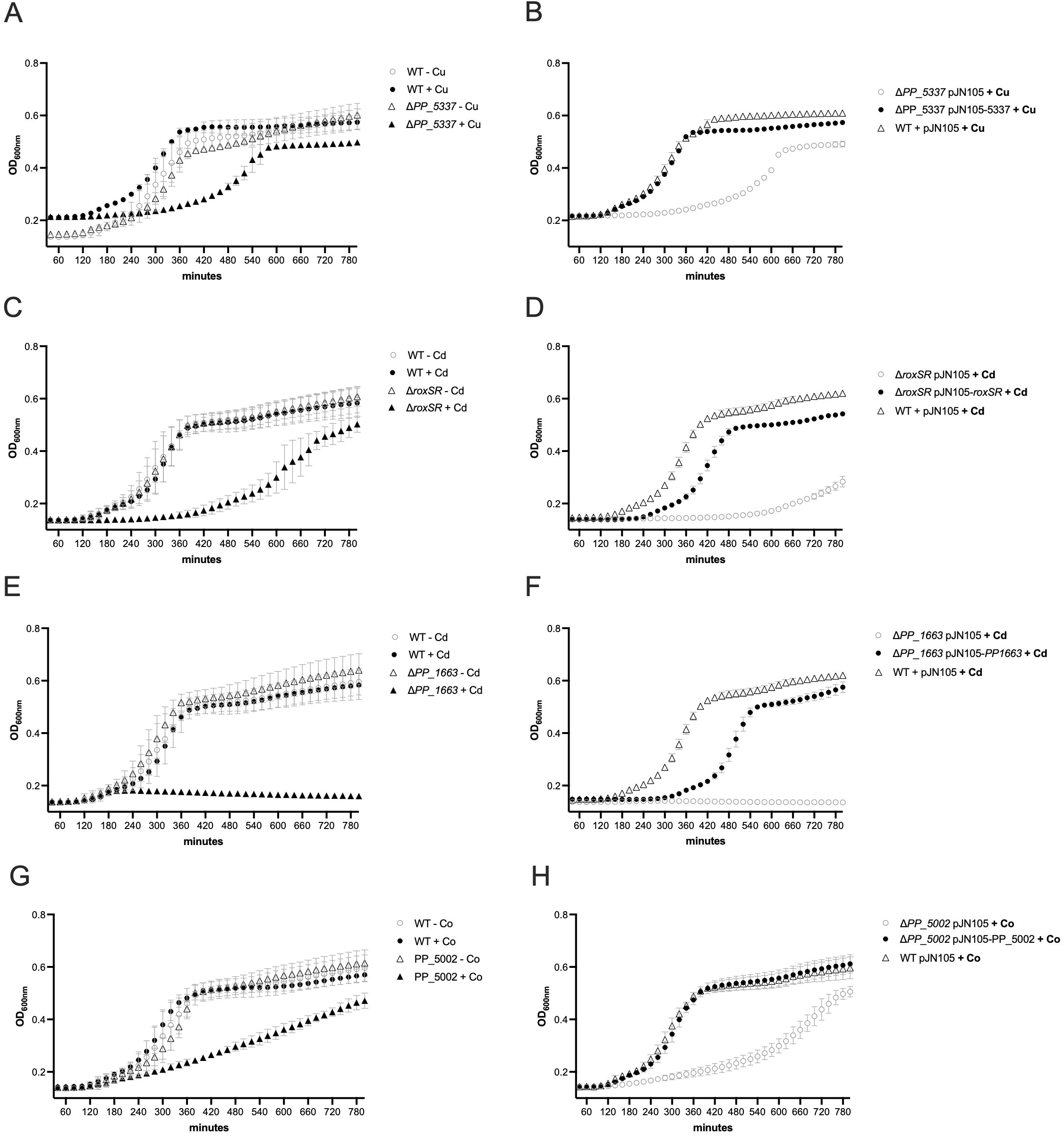
Individual growth cultures of the mutants and complementation assay. Individual growth of each mutant strain was performed in LB medium supplemented or not with a sub-inhibitory concentration of metals (cobalt 10 µM, zinc 125 µM, copper 2.5 mM, cadmium 12.5 µM) in a 96 well plate. OD at _600nm_ was measured over the time. Panels A, C, E and F shows growth of the WT and the mutants in both conditions. Panels B, D, F and H shows the functional complementation assay in presence of metal. The data represents the mean of 4 replicates. The growth difference between the mutant and the WT (panels A, C, E and F) or the complemented mutant and the WT in the presence of a metal (panels B, D, F and H) is always statistically significantly different during the exponential phase (p<0.05, Mann-Whitney U test).

To prove that the phenotypes of the mutants were in fact related to the deletion of the respective target gene, we cloned the genes into the pJN105 plasmid under the control of the arabinose-inducible promoter pBAD to perform a complementation assay. The WT and mutants were grown in LB with arabinose to induce gene expression from the pJN105 plasmid. Cultures were performed in the presence of Cu, Cd, or Co (Fig. 3B, D, F, G). Expression of the *PP_5337, roxSR, PP_1663* and *PP_5002* genes in the corresponding mutant could at least partially suppress the growth defect caused by the metals. In particular, these data demonstrated that *PP_1663* and *roxSR* are novel genetic factors required for Cd^2+^, *PP_5337* for Cu^2+^ and *PP_5002* for Co^2+^ metal resistance in *P. putida* KT2440. Taken together, these results confirm that Tn-seq is (i) a reliable technique for identifying genes involved in metal resistance in *P. putida* and ii) is able to confirm known genes and also identify novel genes relevant for metal resistance.

## Concluding remarks

Tn-seq has previously been used to comprehensively study the essential genomes of several bacteria, sometimes in response to drugs (22, 66). However, there has been no Tn-seq genome-wide study of factors necessary for Cu, Cd, Co and Zn resistance in *Pseudomonas putida*. Motivated by previous studies, we ensured that we could rely on a completely assembled, full genome sequence of our *P. putida* KT2440 strain in order to minimize the risk to miss relevant genes (67). Overall, our approach was risky because *P. putida* has multiple genetic determinants that affect its resistance to these metals. Functional redundancy can be a challenge in this type of experimental approach. The absence of a genetic determinant for metal-resistance may be compensated for by the expression of other metal-resistant genes. We chose to work with sub-inhibitory concentrations of Cu, Cd, Co and Zn. Our approach let to the identification of genes already known to be implicated in metal resistance or homeostasis. The study has identified key genes involved in resistance, such as *copA-1*, *pcoA-2*, and *pcoB-2* for copper, *cadA-3* and *cadR* for cadmium, and *czcA-1* for zinc. This finding is also significant because it indicates that these genes do not have functional redundancy. Miller and colleagues have demonstrated the response of P. putida to the presence of cadmium and copper (7). Numerous transcriptional regulators, outer and inner membrane proteins that form efflux channels and pumps, periplasmic proteins, and stress-related proteins are involved. It is plausible that the genes identified in our screens are part of the initial defense against these metals. If higher concentrations occur, other genes not identified in our screens will come into play. To exemplify this vision, it is worth mentioning the research conducted by Peng and his colleagues (11). They performed RNA-seq on *P. putida* KT2440 grown under varying zinc concentrations. The authors found that the transcriptome of *P. putida* was dependent on the concentration of zinc in the medium. Specifically, at the lowest concentration tested (200 µM in a semi-synthetic medium), only PP_5139 and PP_0043 were overexpressed. These results support our observation that the two genes are necessary for resistance to a zinc concentration of 125 µM in LB.

Another example is the RND complex CzcCBA (PP_0043-PP_0045), known for its resistance to zinc and cadmium (4). It appears that this system is dispensable for cadmium resistance in our screen. It is possible that the *czcABC* system responds to a higher quantity of metals and was therefore inactive in our conditions. This hypothesis is supported by the identification of the *czCBA1* genes involved in resistance to 3mM cadmium resistance during the screening of a Tn5 mutant library in *P. putida* CD2 (12). The difference in the experiments in the two studies could explain this phenomenon. It would thus be interesting to carry out a Tn-seq screen at higher metal concentrations. This would aid in identifying additional genetic factors required for survival in environments with high levels of heavy metals. Tn-seq has the added advantage of being able to identify genes that are not induced in the presence of metal, making its results different from those obtained by transcriptomics or proteomics.

Finally, our study allowed the identification of new important factors for metal resistance in *P. putida*. Targeted in-frame mutagenesis and functional complementation prove that *PP_1663* encoding a periplasmic protein and *roxSR* encoding a two-component system are required for cadmium resistance. *PP_5337* is a new putative transcriptional regulator required for copper resistance, and PP_5002 an hypothetical protein required for cobalt resistance. To better understand how these genes induces resistance to metals, further characterization is necessary. For RoxR and PP_5337, a transcriptomic study should be conducted to identify the genes that are regulated by these transcriptional regulators. In conclusion, our study shows that there are still many studies to be carried out to fully understand the *P. putida* resistome to metals.

## Methods

### Bacterial strains and growth conditions

Bacterial strains, plasmids and oligonucleotides used in this study are described in Table S1 and Table S2. During the course of the project, we decided to re-sequence the genome of our *P. putida* KT2440 strain present in our collection, referred to as PP1 (see supplementary methods). The genome is registered under Genbank accession CP036494. The Average Nucleotide Identity between this strain and *P. putida* KT2440 (Genbank AE015451.2) is 100% (http://enve-omics.ce.gatech.edu/ani/). The PP1 is thus referred as KT2440 strain in the article.

*P. putida* and *E. coli* cells were grown at 28 and 37°C respectively in LB medium or 2YT medium. When required, antibiotics were added at the following concentration: ampicillin, 100 µg/L, gentamicin, 30 µg/L for *P. putida* and 7 µg/L for *E. coli*, streptomycin, 100 µg/L. Media were solidified with 1.5 g/L agar. During Tn-seq experiments, metals were used at a subinhibitory concentration: CoCl_2_ 10 µM, ZnCl_2_ 125 µM, CuCl_2_ 2.5 mM, CdCl_2_ 12.5 µM.

### Construction of the transposon library

*P. putida* strain KT2440 and *E. coli* MFDpir/pEGL55 were grown overnight in 2YT medium. pEGL55 is a R6K suicide plasmid carrying the mariner transposon. 100 OD_600nm_ units of each strain were mixed and centrifuged at 5000g for 10 min. The bacteria were resuspended in 1.2 mL of 2YT medium supplemented with diaminopimelic acid (300 µM) and plated on an over-dried LB agar plate containing twice the normal concentration of agar. After 3 hours at 28°C, bacteria were collected and resuspended in 4 mL LB medium. A 20 µl aliquot was diluted and plated on LB agar with gentamicin to estimate the efficiency of mutagenesis. The other part was spread on 50 plates of LB agar with gentamicin and grown for 24 h at 28°C. To confirm that the *P. putida* mutants had lost the plasmid, we performed colony PCR with primers annealing to the *bla* gene of pEGL55. None of the 100 colonies tested produced a PCR fragment, indicating loss of the plasmid in the bacteria tested. 800,000 mutants were harvested in LB supplemented with 40% glycerol at −80°C. This library was directly sequenced and represents the mutant pool in LB agar (see Table S4).

### DNA preparation for high-throughput sequencing

To identify essential genes in LB or LB with metal, ∼ 10^7^ mutants were inoculated in 25 mL LB. The culture was then incubated at 28°C with shaking at 180 rpm. At OD_600_ of 0.2, metals were added independently at the following subinhibitory concentrations: cobalt 10 µM, copper 2.5 mM, zinc 125 µM, and cadmium 12.5 µM. When OD_600_ was 1.6, the culture medium was diluted in the same medium with OD_600_ of 0.03. This procedure was carried out for 12 generations. The final pools of mutants were harvested by centrifugation of the culture medium and stored at −80°C. DNA was extracted from aliquots of the bacterial suspension using the Promega Wizard Genomic DNA Purification Kit. The next steps of the DNA preparation methods were performed as described previously (20). Quality control of Tn-seq DNA libraries (fragment size and concentration) and high-throughput sequencing on HiSeq 2500 (Illumina) were performed by MGX (CNRS sequencing service, Montpellier). 6 DNA libraries were multiplexed on a flow cell. After demultiplexing, the total number of reads ranged from 19 to 35 million (Table S3).

### Bioinformatics analysis

Raw reads from the fastQ files were first filtered using cutadapt v1.11 (Martin, 2011), and only reads containing the mariner inverted left repeat (ACAGGTTGGATGATAAGTCCCCGGTCTT) were trimmed and considered bona fide transposon disrupted genes. The trimmed reads were then analyzed using a modified version of the TPP script available in the TRANSIT software v2.0.2 (26447887). The mapping step was modified to select only reads that mapped uniquely and without mismatch in the P. putida KT2440 genome. The counting step was then modified to accurately count reads mapping to each TA site in the reference genome according to the Tn-seq protocol used in this study. Read counts per insertion were normalized using the LOESS method as described in Zomer et al. (68). Next, we used the TRANSIT software (version 2.0) to compare the Tn-seq datasets (27). Gene states obtained by TRANSIT after growth of the mutant bank of *P. putida* KT24440 in LB agar and LB are presented in Table S4. Raw data of all datasets analyzed by TRANSIT are presented in Table S5.

#### Construction of the pKNG101 plasmids used for in-frame deletion in *P. putida* (Table S1)

The 500 bp of DNA upstream and downstream of a target gene were amplified by PCR (Primestar Max DNA Polymerase, Takara). The two 500 bp fragments were then fused by overlapping PCR. The resulting 1 kbp DNA fragment was inserted between the BamHI/SpeI restriction sites of pKNG101 by SLIC (69). Finally, the construct was transformed into DH5□ □pir and verified by colony PCR and sequencing.

#### Construction of the pJN105 plasmids used for complementation (Table S2)

The target gene with native RBS was amplified by PCR (Primestar Max DNA Polymerase) from gDNA of P. putida KT2440. The amplified fragment was inserted by SLIC between the SpeI and SacI restriction sites of pJN105 and then transformed into DH5L. The resulting plasmids were validated by restriction mapping and sequencing.

### In-frame deletion mutant construction

To construct the in-frame deletion mutants of the genes underlined in Figure 2, the counter selection method using the sacB gene was used (70). The suicide pKNG101 plasmid were transferred from MFD*pir* (71) to *P. putida* KT2440. The first recombination event was selected on LB agar supplemented with streptomycin. Transconjugants were then plated on LB agar without NaCl supplemented with 5% sucrose to allow the second recombination event. In-frame deletions were then verified by PCR (Dreamtaq polymerase, Thermofisher).

### 1 × 1 Competition assays

To compare the metal sensitivity of the mutants with the wild-type strain, 1 × 1 competition experiments were performed as follows. First, to distinguish the mutants from the wild strain, a GFP^+^ WT strain was constructed by inserting the constitutively expressed gfp gene into the attTn7 site of the *P. putida* KT2440 chromosome using the pUC18-miniTn7-gfpmut3 plasmid (72). The GFP^+^ strain grow as well as the WT (figure S4). Mutant and GFP^+^ WT strains were grown separately in LB medium from an overnight culture in LB to OD_600_ of 0.8. Bacteria were then mixed in a 1:1 ratio at an initial OD_600_ of 0.0125 in a 96-well plate containing 200 µL LB or LB with metal at a sub-inhibitory concentration. After 24 hours of growth at 28°C in the Tecan M200 Pro with shaking, 5 µL of the cultures were used to inoculate a new 96-well plate and placed under the same conditions. After a total of 48 hours of growth (approximately 10 divisions), the bacteria were diluted and plated onto LB agar plates. After 48 hours at 28°C, GFP^+^ wild-type and mutant colonies were counted under blue light to detect colony fluorescence. A ratio was then calculated by dividing the number of mutant colonies by the number of wild-type colonies in each condition. The growth comparison between a WT strain and a GFP^+^ strain in LB supplemented with different metals is shown in Table S6.

### Individual growth in presence of metals

Single strain growth was performed in LB medium from an overnight culture in LB to an OD_600_ of 0.8. Bacteria were then inoculated at an initial OD_600_ of 0.006 into a 96-well plate containing 200 µL of LB or LB with metal at a sub-inhibitory concentration and placed at 28°C in the Tecan M200 Pro. OD_600_ measurements were taken every 10 minutes after shaking. Complementation assays were performed using the same protocol but with 0.2% arabinose. Data are presented after 6.5 hours of growth.

## Acknowledgment

We thank Geraldine Effantin, Veronique Utzinger for technical assistance, the members of the MTSB team and Xavier Charpentier, Bérengère Ize, Sylvie Elsen, for discussion.

## References

1. Nelson KE, Weinel C, Paulsen IT, Dodson RJ, Hilbert H, Martins dos Santos VAP, Fouts DE, Gill SR, Pop M, Holmes M, Brinkac L, Beanan M, DeBoy RT, Daugherty S, Kolonay J, Madupu R, Nelson W, White O, Peterson J, Khouri H, Hance I, Chris Lee P, Holtzapple E, Scanlan D, Tran K, Moazzez A, Utterback T, Rizzo M, Lee K, Kosack D, Moestl D, Wedler H, Lauber J, Stjepandic D, Hoheisel J, Straetz M, Heim S, Kiewitz C, Eisen JA, Timmis KN, Düsterhöft A, Tümmler B, Fraser CM. 2002. Complete genome sequence and comparative analysis of the metabolically versatile Pseudomonas putida KT2440. Environ Microbiol 4:799–808.

2. Wu X, Monchy S, Taghavi S, Zhu W, Ramos J, van der Lelie D. 2011. Comparative genomics and functional analysis of niche-specific adaptation in Pseudomonas putida. FEMS Microbiol Rev 35:299–323.

3. Clarke PH. 1982. The metabolic versatility of pseudomonads. Antonie Van Leeuwenhoek 48:105–130.

4. Cánovas D, Cases I, de Lorenzo V. 2003. Heavy metal tolerance and metal homeostasis in Pseudomonas putida as revealed by complete genome analysis. Environ Microbiol 5:1242–1256.

5. Bruins MR, Kapil S, Oehme FW. 2000. Microbial resistance to metals in the environment. Ecotoxicol Environ Saf 45:198–207.

6. Chandrangsu P, Rensing C, Helmann JD. 2017. Metal homeostasis and resistance in bacteria. Nat Rev Microbiol 1–13.

7. Miller CD, Pettee B, Zhang C, Pabst M, McLean JE, Anderson AJ. 2009. Copper and cadmium: responses in Pseudomonas putida KT2440. Lett Appl Microbiol 49:775–783.

8. Manara A, DalCorso G, Baliardini C, Farinati S, Cecconi D, Furini A. 2012. Pseudomonas putida response to cadmium: changes in membrane and cytosolic proteomes. J Proteome Res 11:4169–4179.

9. Cheng Z, Wei Y-YC, Sung WWL, Glick BR, McConkey BJ. 2009. Proteomic analysis of the response of the plant growth-promoting bacterium Pseudomonas putida UW4 to nickel stress. Proteome Sci 7:18.

10. Ray P, Girard V, Gault M, Job C, Bonneu M, Mandrand-Berthelot M-A, Singh SS, Job D, Rodrigue A. 2013. Pseudomonas putida KT2440 response to nickel or cobalt induced stress by quantitative proteomics. Met Integr Biometal Sci 5:68–79.

11. Peng J, Miao L, Chen X, Liu P. 2018. Comparative Transcriptome Analysis of Pseudomonas putida KT2440 Revealed Its Response Mechanisms to Elevated Levels of Zinc Stress. Front Microbiol 9.

12. Hu N, Zhao B. 2007. Key genes involved in heavy-metal resistance in Pseudomonas putida CD2. FEMS Microbiol Lett.

13. Molina-Henares MA, De La Torre J, García-Salamanca A, Molina-Henares AJ, Herrera MC, Ramos JL, Duque E. 2010. Identification of conditionally essential genes for growth of Pseudomonas putida KT2440 on minimal medium through the screening of a genome-wide mutant library. Environ Microbiol 12:1468–1485.

14. van Opijnen T, Bodi KL, Camilli A. 2009. Tn-seq: high-throughput parallel sequencing for fitness and genetic interaction studies in microorganisms. Nat Methods 6:767–772.

15. van Opijnen T, Levin HL. 2020. Transposon Insertion Sequencing, a Global Measure of Gene Function. Annu Rev Genet 54:337–365.

16. Goodman AL, McNulty NP, Zhao Y, Leip D, Mitra RD, Lozupone CA, Knight R, Gordon JI. 2009. Identifying genetic determinants needed to establish a human gut symbiont in its habitat. Cell Host Microbe 6:279–289.

17. Fu Y, Waldor MK, Mekalanos JJ. 2013. Tn-Seq analysis of vibrio cholerae intestinal colonization reveals a role for T6SS-mediated antibacterial activity in the host. Cell Host Microbe 14:652–663.

18. Skurnik D, Roux D, Aschard H, Cattoir V, Yoder-Himes D, Lory S, Pier GB. 2013. A Comprehensive Analysis of In Vitro and In Vivo Genetic Fitness of Pseudomonas aeruginosa Using HighThroughput Sequencing of Transposon Libraries. PLoS Pathog 9:e1003582.

19. Helmann TC, Deutschbauer AM, Lindow SE. 2019. Genome-wide identification of Pseudomonas syringae genes required for fitness during colonization of the leaf surface and apoplast. Proc Natl Acad Sci U S A 116:18900–18910.

20. Royet K, Parisot N, Rodrigue A, Gueguen E, Condemine G. 2019. Identification by Tn-seq of Dickeya dadantii genes required for survival in chicory plants. Mol Plant Pathol 20:287–306.

21. Morinière L, Mirabel L, Gueguen E, Bertolla F. 2022. A Comprehensive Overview of the Genes and Functions Required for Lettuce Infection by the Hemibiotrophic Phytopathogen Xanthomonas hortorum pv. vitians. mSystems 7:e0129021.

22. Gallagher LA, Shendure J, Manoil C. 2011. Genome-scale identification of resistance functions in Pseudomonas aeruginosa using Tn-seq. mBio 2:e00315–10.

23. Calero P, Jensen SI, Bojanovič K, Lennen R, Koza A, Nielsen AT. 2017. Genome-wide identification of tolerance mechanisms towards p-coumaric acid in Pseudomonas putida. Biotechnol Bioeng 10.1002/bit.26495.

24. Borchert AJ, Bleem A, Beckham GT. 2023. RB-TnSeq identifies genetic targets for improved tolerance of *Pseudomonas putida* towards compounds relevant to lignin conversion. Metab Eng 77:208–218.

25. Higgins S, Gualdi S, PintoLCarbó M, Eberl L. 2020. Copper resistance genes of Burkholderia cenocepacia H111 identified by transposon sequencing. Environ Microbiol Rep 12:241–249.

26. Gualdi S, Agnoli K, Vitale A, Higgins S, Eberl L. 2022. Identification of genes required for gold and silver tolerance in Burkholderia cenocepacia H111 by transposon sequencing. Environ Microbiol 24:737–751.

27. DeJesus MA, Ambadipudi C, Baker R, Sassetti C, Ioerger TR. 2015. TRANSIT--A Software Tool for Himar1 TnSeq Analysis. PLoS Comput Biol 11:e1004401.

28. Teitzel GM, Geddie A, De Long SK, Kirisits MJ, Whiteley M, Parsek MR. 2006. Survival and growth in the presence of elevated copper: transcriptional profiling of copper-stressed Pseudomonas aeruginosa. J Bacteriol 188:7242–7256.

29. Thaden JT, Lory S, Gardner TS. 2010. Quorum-Sensing Regulation of a Copper Toxicity System in Pseudomonas aeruginosa. J Bacteriol 192:2557–2568.

30. Quintana J, Novoa-Aponte L, Argüello JM. 2017. Copper homeostasis networks in the bacterium Pseudomonas aeruginosa. J Biol Chem 292:15691–15704.

31. González-Guerrero M, Raimunda D, Cheng X, Argüello JM. 2010. Distinct functional roles of homologous Cu+ efflux ATPases in Pseudomonas aeruginosa. Mol Microbiol 78:1246–1258.

32. Varadi M, Anyango S, Deshpande M, Nair S, Natassia C, Yordanova G, Yuan D, Stroe O, Wood G, Laydon A, Žídek A, Green T, Tunyasuvunakool K, Petersen S, Jumper J, Clancy E, Green R, Vora A, Lutfi M, Figurnov M, Cowie A, Hobbs N, Kohli P, Kleywegt G, Birney E, Hassabis D, Velankar S. 2022. AlphaFold Protein Structure Database: massively expanding the structural coverage of protein-sequence space with high-accuracy models. Nucleic Acids Res 50:D439–D444.

33. Zhang Y, Skolnick J. 2005. TM-align: a protein structure alignment algorithm based on the TM-score. Nucleic Acids Res 33:2302–2309.

34. Novichkov PS, Kazakov AE, Ravcheev DA, Leyn SA, Kovaleva GY, Sutormin RA, Kazanov MD, Riehl W, Arkin AP, Dubchak I, Rodionov DA. 2013. RegPrecise 3.0--a resource for genome-scale exploration of transcriptional regulation in bacteria. BMC Genomics 14:745.

35. Lee SM, Grass G, Rensing C, Barrett SR, Yates CJD, Stoyanov JV, Brown NL. 2002. The Pco proteins are involved in periplasmic copper handling in *Escherichia coli*. Biochem Biophys Res Commun 295:616–620.

36. Cha JS, Cooksey DA. 1991. Copper resistance in Pseudomonas syringae mediated by periplasmic and outer membrane proteins. Proc Natl Acad Sci U S A 88:8915–8919.

37. Outten FW, Huffman DL, Hale JA, O’Halloran TV. 2001. The Independent cue and cusSystems Confer Copper Tolerance during Aerobic and Anaerobic Growth inEscherichia coli *. J Biol Chem 276:30670–30677.

38. Gudipaty SA, Larsen AS, Rensing C, McEvoy MM. 2012. Regulation of Cu(I)/Ag(I) efflux genes in Escherichia coli by the sensor kinase CusS. FEMS Microbiol Lett 330:30–37.

39. Leedjärv A, Ivask A, Virta M. 2008. Interplay of different transporters in the mediation of divalent heavy metal resistance in Pseudomonas putida KT2440. J Bacteriol 190:2680–2689.

40. Nucifora G, Chu L, Misra TK, Silver S. 1989. Cadmium Resistance from Staphylococcus aureus Plasmid pI258 cadA Gene Results from a Cadmium-Efflux ATPase. Proc Natl Acad Sci U S A 86:3544–3548.

41. Rensing C, Mitra B, Rosen BP. 1997. The zntA gene of Escherichia coli encodes a Zn(II)-translocating P-typeLATPase. Proc Natl Acad Sci 94:14326–14331.

42. Ito K, Inaba K. 2008. The disulfide bond formation (Dsb) system. Curr Opin Struct Biol 18:450–458.

43. Rensing C, Mitra B, Rosen BP. 1997. Insertional inactivation of dsbA produces sensitivity to cadmium and zinc in Escherichia coli. J Bacteriol 179:2769–2771.

44. Hayashi S, Abe M, Kimoto M, Furukawa S, Nakazawa T. 2000. The dsbA-dsbB disulfide bond formation system of Burkholderia cepacia is involved in the production of protease and alkaline phosphatase, motility, metal resistance, and multi-drug resistance. Microbiol Immunol 44:41–50.

45. Fernández-Piñar R, Ramos JL, Rodríguez-Herva JJ, Espinosa-Urgel M. 2008. A two-component regulatory system integrates redox state and population density sensing in Pseudomonas putida. J Bacteriol 190:7666–7674.

46. Nies DH. 2003. Efflux-mediated heavy metal resistance in prokaryotes. FEMS Microbiol Rev 27:313–339.

47. Nies DH, Rehbein G, Hoffmann T, Baumann C, Grosse C. 2006. Paralogs of Genes Encoding Metal Resistance Proteins in Cupriavidus metallidurans Strain CH34. J Mol Microbiol Biotechnol 11:82–93.

48. Anton A, Große C, Reißmann J, Pribyl T, Nies DH. 1999. CzcD Is a Heavy Metal Ion Transporter Involved in Regulation of Heavy Metal Resistance in Ralstonia sp. Strain CH34. J Bacteriol 181:6876–6881.

49. Haritha A, Sagar KP, Tiwari A, Kiranmayi P, Rodrigue A, Mohan PM, Singh SS. 2009. MrdH, a novel metal resistance determinant of Pseudomonas putida KT2440, is flanked by metal-inducible mobile genetic elements. J Bacteriol 191:5976–5987.

50. Snavely MD, Florer JB, Miller CG, Maguire ME. 1989. Magnesium transport in Salmonella typhimurium: 28Mg2+ transport by the CorA, MgtA, and MgtB systems. J Bacteriol 171:4761–4766.

51. Smith RL, Maguire ME. 1998. Microbial magnesium transport: unusual transporters searching for identity. Mol Microbiol 28:217–226.

52. Blencowe DK, Morby AP. 2003. Zn(II) metabolism in prokaryotes. FEMS Microbiol Rev 27:291–311.

53. Hantke K. 2005. Bacterial zinc uptake and regulators. Curr Opin Microbiol 8:196–202.

54. Schalk IJ, Guillon L. 2013. Pyoverdine biosynthesis and secretion in *P seudomonas aeruginosa* L: implications for metal homeostasis. Environ Microbiol 15:1661–1673.

55. Ringel MT, Dräger G, Brüser T. 2016. PvdN Enzyme Catalyzes a Periplasmic Pyoverdine Modification. J Biol Chem 291:23929–23938.

56. Ringel MT, Dräger G, Brüser T. 2018. PvdO is required for the oxidation of dihydropyoverdine as the last step of fluorophore formation in Pseudomonas fluorescens. J Biol Chem 293:2330–2341.

57. Sugue M-F, Burdur AN, Ringel MT, Dräger G, Brüser T. 2022. PvdM of fluorescent pseudomonads is required for the oxidation of ferribactin by PvdP in periplasmic pyoverdine maturation. J Biol Chem 298.

58. Braud A, Geoffroy V, Hoegy F, Mislin GLA, Schalk IJ. 2010. Presence of the siderophores pyoverdine and pyochelin in the extracellular medium reduces toxic metal accumulation in Pseudomonas aeruginosa and increases bacterial metal tolerance. Environ Microbiol Rep 2:419–425.

59. Helbig K, Bleuel C, Krauss GJ, Nies DH. 2008. Glutathione and transition-metal homeostasis in Escherichia coli. J Bacteriol 190:5431–5438.

60. Stewart LJ, Ong CY, Zhang MM, Brouwer S, McIntyre L, Davies MR, Walker MJ, McEwan AG, Waldron KJ, Djoko KY. 2020. Role of Glutathione in Buffering Excess Intracellular Copper in Streptococcus pyogenes. mBio 11:10.1128/mbio.02804-20.

61. Lee S-W, Glickmann E, Cooksey DA. 2001. Chromosomal Locus for Cadmium Resistance in Pseudomonas putida Consisting of a Cadmium-Transporting ATPase and a MerR Family Response Regulator. Appl Environ Microbiol 67:1437–1444.

62. Liu X, Hu Q, Yang J, Huang S, Wei T, Chen W, He Y, Wang D, Liu Z, Wang K, Gan J, Chen H. 2019. Selective cadmium regulation mediated by a cooperative binding mechanism in CadR. Proc Natl Acad Sci 116:20398–20403.

63. Ducret V, Gonzalez MR, Leoni S, Valentini M, Perron K. 2020. The CzcCBA Efflux System Requires the CadA P-Type ATPase for Timely Expression Upon Zinc Excess in Pseudomonas aeruginosa. Front Microbiol 11:911.

64. Seo J, Brencic A, Darwin AJ. 2009. Analysis of secretin-induced stress in Pseudomonas aeruginosa suggests prevention rather than response and identifies a novel protein involved in secretin function. J Bacteriol 191:898–908.

65. Putker F, Tommassen-van Boxtel R, Stork M, Rodríguez-Herva JJ, Koster M, Tommassen J. 2013. The type II secretion system (Xcp) of Pseudomonas putida is active and involved in the secretion of phosphatases. Environ Microbiol 15:2658–2671.

66. Barquist L, Boinett CJ, Cain AK. 2013. Approaches to querying bacterial genomes with transposon-insertion sequencing. RNA Biol 10.

67. Varadarajan AR, Allan RN, Valentin JDP, Castañeda Ocampo OE, Somerville V, Pietsch F, Buhmann MT, West J, Skipp PJ, van der Mei HC, Ren Q, Schreiber F, Webb JS, Ahrens CH. 2020. An integrated model system to gain mechanistic insights into biofilm-associated antimicrobial resistance in Pseudomonas aeruginosa MPAO1. NPJ Biofilms Microbiomes 6:46.

68. Zomer A, Burghout P, Bootsma HJ, Hermans PWM, van Hijum SAFT. 2012. ESSENTIALS: software for rapid analysis of high throughput transposon insertion sequencing data. PLoS ONE 7:e43012.

69. Jeong J-Y, Yim H-S, Ryu J-Y, Lee HS, Lee J-H, Seen D-S, Kang SG. 2012. One-step sequence- and ligation-independent cloning as a rapid and versatile cloning method for functional genomics studies. Appl Environ Microbiol 78:5440–5443.

70. Kaniga K, Delor I, Cornelis GR. 1991. A wide-host-range suicide vector for improving reverse genetics in gram-negative bacteria: inactivation of the blaA gene of Yersinia enterocolitica. Gene 109:137–141.

71. Ferrières L, Hémery G, Nham T, Guérout A-M, Mazel D, Beloin C, Ghigo J-M. 2010. Silent Mischief: Bacteriophage Mu Insertions Contaminate Products of Escherichia coli Random Mutagenesis Performed Using Suicidal Transposon Delivery Plasmids Mobilized by Broad-Host-Range RP4 Conjugative Machinery. J Bacteriol 192:6418–6427.

72. Choi K-H, Schweizer HP. 2006. mini-Tn7 insertion in bacteria with single attTn7 sites: example Pseudomonas aeruginosa. Nat Protoc 1:153–161.

73. Winsor GL, Griffiths EJ, Lo R, Dhillon BK, Shay JA, Brinkman FSL. 2016. Enhanced annotations and features for comparing thousands of Pseudomonas genomes in the Pseudomonas genome database. Nucleic Acids Res 44:D646–D653.

74. The UniProt Consortium. 2015. UniProt: a hub for protein information. Nucleic Acids Res 43:D204–D212.

