## Supplementary figures and images for "High-throughput Tn-seq screens identify both known and novel *Pseudomonas putida* KT2440 genes involved in metal resistance"

### fig. S1

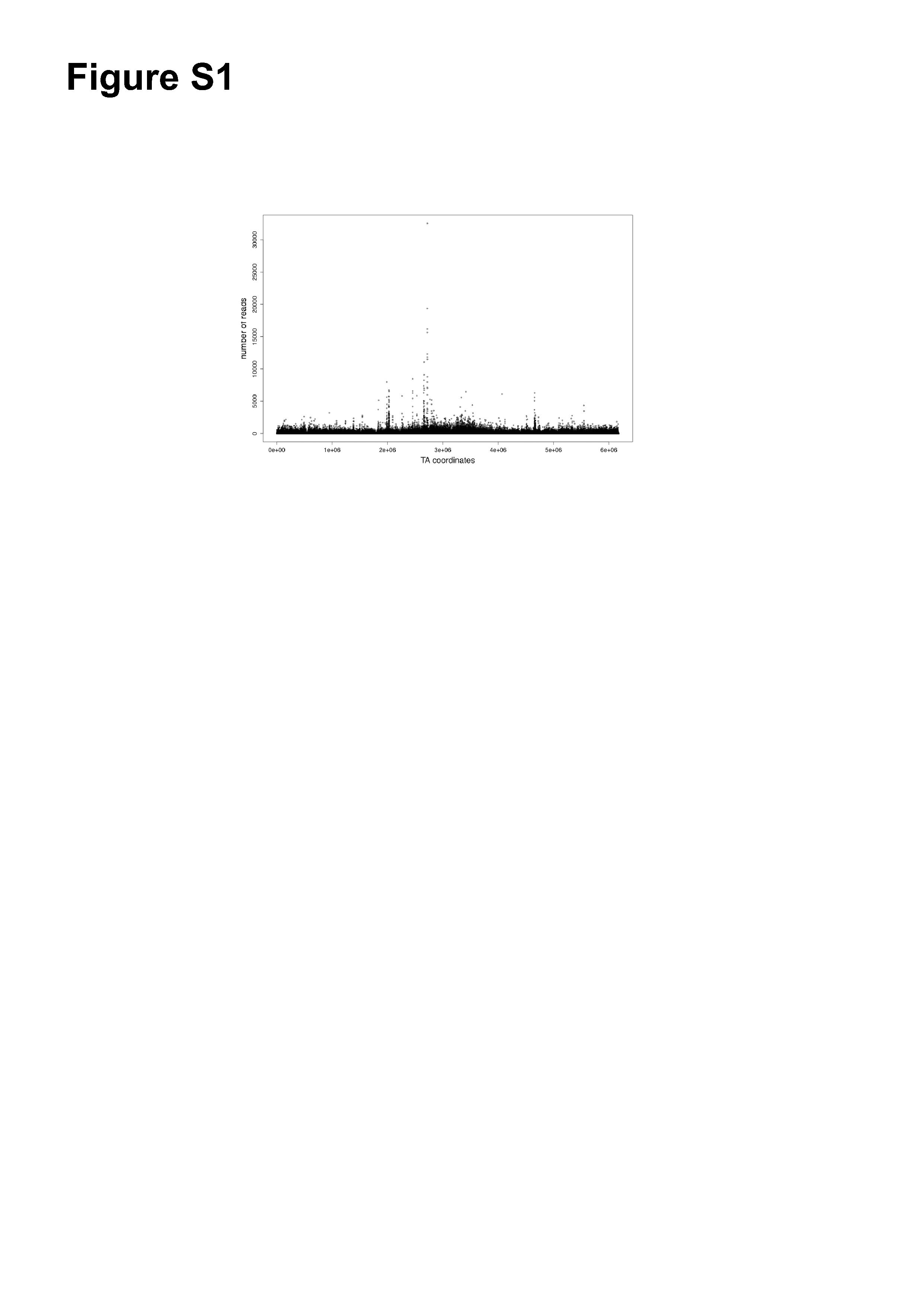

### fig. S2

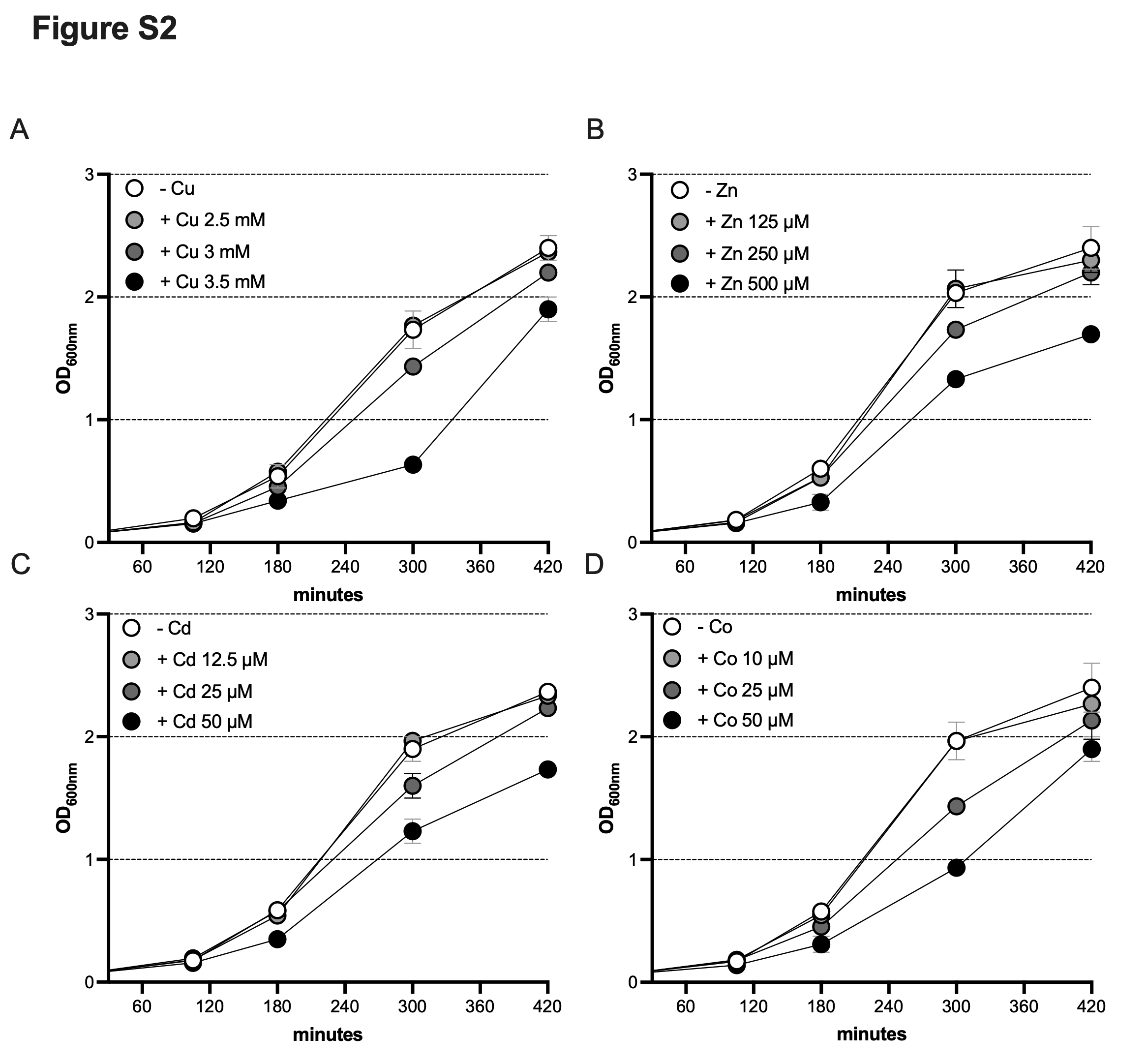

### fig. S3

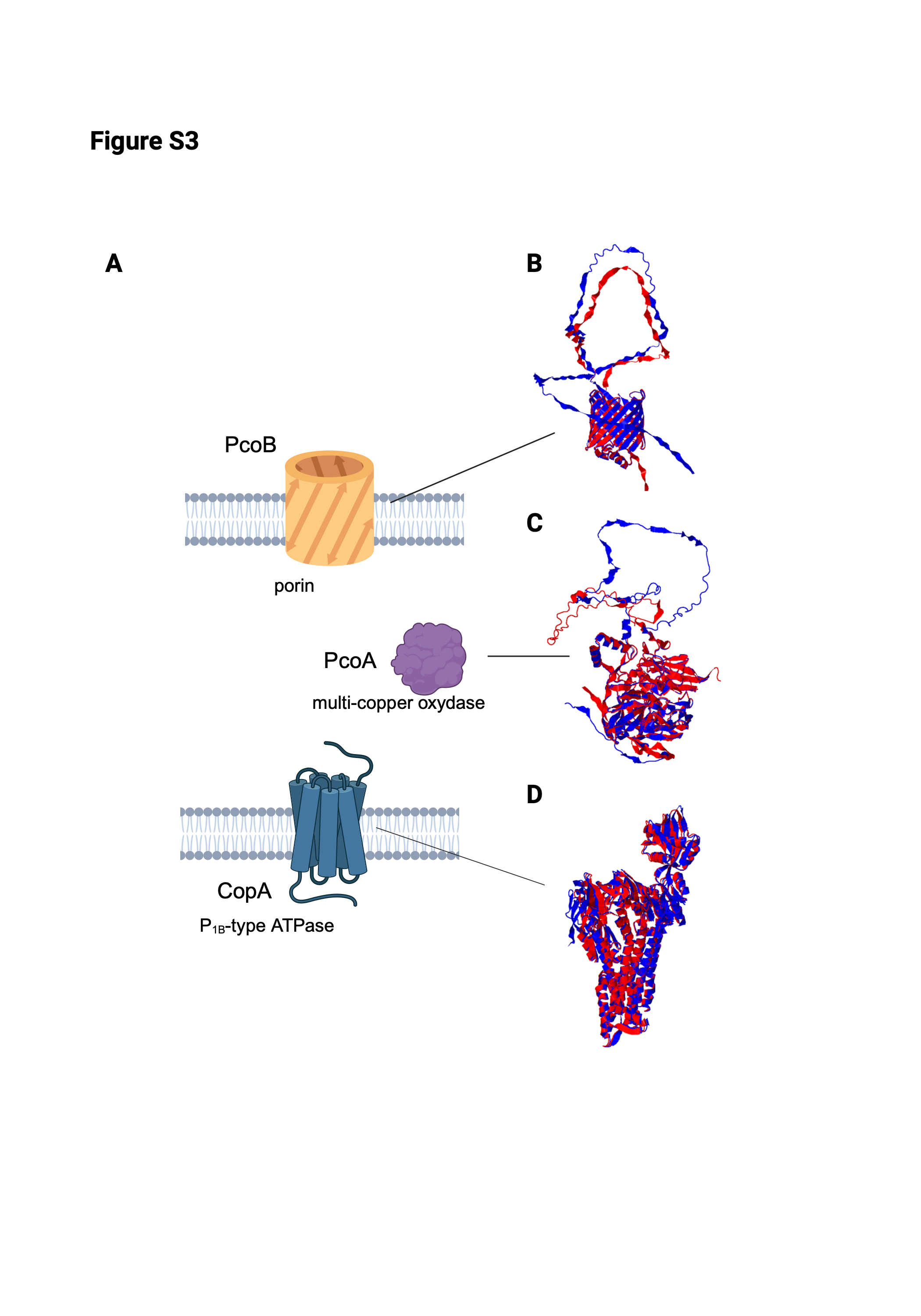

### fig. S4

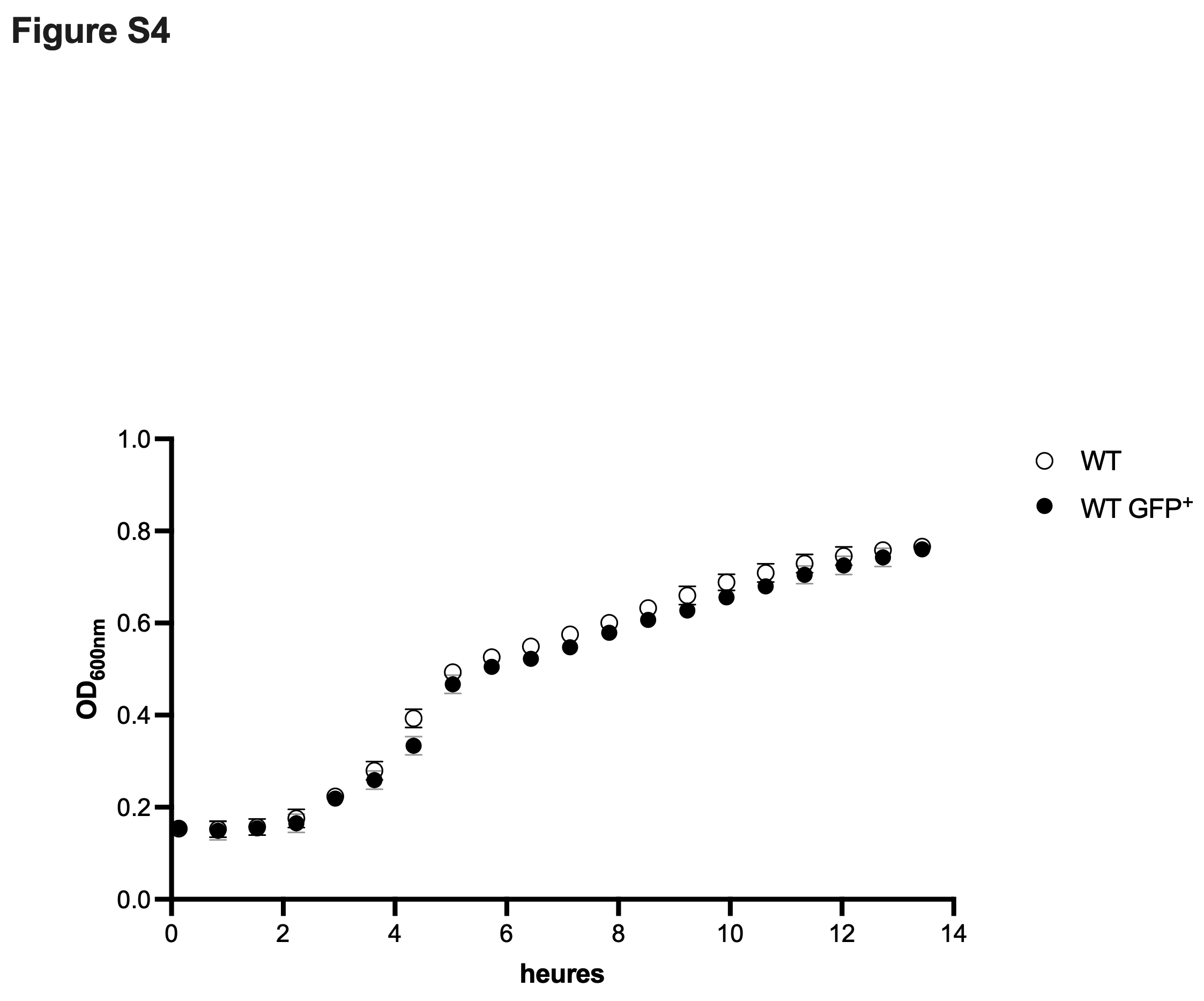
