## Supplementary methods for "High-throughput Tn-seq screens identify both known and novel *Pseudomonas putida* KT2440 genes involved in metal resistance"

**Sequencing, *de novo* genome assembly and annotation of the Pseudomonas putida PP1 strain**

gDNA of the PP1 strain was extracted using the GenElute kit (Sigma). PacBio SMRT sequencing was carried out on an RSII machine (1 SMRT cell per strain). Size selection was performed using the BluePippin system and resulted in fragments with an average subread length of 17-20 kb. PacBio RSII subreads were extracted using the SMRT Portal and protocol “RS_HGAP_Assembly.3” (default parameters, except: minimum subread length of 5000, estimated genome size of 6 Mb).

Two 2 x 300 bp Illumina paired end libraries were prepared per sample using the Nextera XT DNA kit and sequenced on a MiSeq. A subset of the highest quality reads (“LEADING:3 TRAILING:3 SLIDINGWINDOW:4:15 MINLEN:36”, rq>15) were extracted using trimmomatic (v. 0.36) (Bolger *et al.*, 2014); only paired reads were used for further analyses.

The filtered PacBio subreads were assembled using Flye (v.2.3.3; default parameters, except: estimated genome size of 6 Mb) (Kolmogorov *et al.*, 2019). All assemblies resulted in one contig each, which were start-aligned using the dnaA gene as start position and polished using 2-5 Quiver runs (Table 3) and the “RS_Resequencing.1” protocol on the SMRT Portal (default parameters, except: minimum subread length of 5000 bp). The genome of *P. putida* KT2440 had to be re-assembled using a minimum subread length of 15 kb in order to resolve a repeat region and was polished using a minimum subread length of 5000 bp and two Quiver runs. To verify the circularity and completeness of the *de novo* assembly, the filtered PacBio subreads were mapped to the circular chromosome using graphmap (v.0.5.2) (Sović *et al.*, 2016). The assembly was further polished using 2 x 300 bp paired end Illumina MiSeq reads and 1-2 Freebayes runs (v.1.2.0; minimum alternate fraction: 0.5, minimum alternate count: 5) (Garrison and Marth, 2012) to correct small errors (e.g., homopolymer errors). Variants were manually inspected in the Integrated Genome Viewer (Thorvaldsdóttir *et al.*, 2013) and subsequently corrected using bcftools (v.0.1.19) (Narasimhan *et al.*, 2016).

The final assemblies were annotated with the NCBI Prokaryotic Genome Annotation Pipeline (Tatusova *et al.*, 2016).

**References**

Bolger, A.M., Lohse, M., and Usadel, B. (2014) Trimmomatic: a flexible trimmer for Illumina sequence data. *Bioinformatics* **30**: 2114–2120.

Garrison, E. and Marth, G. (2012) Haplotype-based variant detection from short-read sequencing.

Kolmogorov, M., Yuan, J., Lin, Y., and Pevzner, P.A. (2019) Assembly of long, error-prone reads using repeat graphs. *Nat Biotechnol* **37**: 540–546.

Narasimhan, V., Danecek, P., Scally, A., Xue, Y., Tyler-Smith, C., and Durbin, R. (2016) BCFtools/RoH: a hidden Markov model approach for detecting autozygosity from next-generation sequencing data. *Bioinformatics* **32**: 1749–1751.

Sović, I., Šikić, M., Wilm, A., Fenlon, S.N., Chen, S., and Nagarajan, N. (2016) Fast and sensitive mapping of nanopore sequencing reads with GraphMap. *Nat Commun* **7**: 11307.

Tatusova, T., DiCuccio, M., Badretdin, A., Chetvernin, V., Nawrocki, E.P., Zaslavsky, L., et al. (2016) NCBI prokaryotic genome annotation pipeline. *Nucleic Acids Res* **44**: 6614–6624.

Thorvaldsdóttir, H., Robinson, J.T., and Mesirov, J.P. (2013) Integrative Genomics Viewer (IGV): high-performance genomics data visualization and exploration. *Brief Bioinform* **14**: 178–192.
